# Dll4 and Jag1a signalling act sequentially and cooperatively to drive hematopoietic stem cell fate specification

**DOI:** 10.64898/2026.08.17.745164

**Authors:** Dashuai Wu, Benjamin Edginton-White, Dorothee Bornhorst, Amulya V. Hejjaji, Felix Gunawan, Rui Monteiro

**Affiliations:** Department of Cancer and Genomic Sciences, School of Medical Sciences, College of Medicine and Health, University of Birmingham, UK; Birmingham Centre for Genome Biology, University of Birmingham, UK; Institute of Cell Biology, Faculty of Medicine, University of Münster, Germany; Institute for Reproductive and Regenerative Biology, Center of Reproductive Medicine and Andrology, Faculty of Medicine, University of Münster, Germany; Department of Cell and Systems Biology, Faculty of Arts and Science, University of Toronto, Canada

## Abstract

Hematopoietic stem and progenitor cells (HSPCs) arise from a specialized subset of arterial endothelial cells, the hemogenic endothelium (HE), during embryonic development and sustain blood production throughout life. Notch signalling is a key regulator of this process: its ligand Jag1 promotes HSPC formation, whereas Dll4 promotes arterial identity. However, how the activities of these ligands are temporally coordinated during HSPC emergence remains unresolved. Here we demonstrate that Dll4 is required prior to circulation onset, acting by dampening MAPK signalling to drive the transition from pre-HE to HE fate and enabling HE differentiation towards HSPCs. Subsequently, after circulation starts, Jag1a acts to maintain gene expression in HE and support transition to HSPC fate. Jag1a activity depends on blood flow-induced shear stress and rescues HSPC loss caused by impaired flow. Thus, rather than playing opposing roles, Dll4 and Jag1a act sequentially and coordinately to drive the endothelial-to-hematopoietic transition and promote HSPC emergence.

## Introduction

Hematopoietic stem and progenitor cells (HSPCs) arise during embryogenesis from a transient population of hemogenic endothelium (HE) located in the ventral wall of the embryonic dorsal aorta (DA), through an endothelial-to-hematopoietic transition (EHT) (Kissa and Herbomel 2010; Boisset et al. 2010; Bertrand et al. 2010). Following EHT, zebrafish HSPCs migrate to the caudal hematopoietic tissue (CHT), a transient hematopoietic niche analogous to the mammalian foetal liver, where they proliferate and differentiate before ultimately colonizing the kidney marrow, the functional equivalent of mammalian bone marrow (Ciau-Uitz et al. 2014). This process is well conserved in vertebrates including humans, mice, and zebrafish (Dzierzak and Bigas 2018). Although EHT is crucial for definitive haematopoiesis and the transcriptional priming state of hemogenic endothelial cells has been characterized (Ghersi et al. 2023), the molecular mechanisms governing the acquisition of hemogenic potential remain incompletely understood.

Notch signalling plays a central role in coordinating arterial identity and hematopoietic specification in the embryonic DA (Lawson et al. 2001; Butko, Pouget, and Traver 2016). The ligands Dll4 and Jag1 have been shown to exert different effects on endothelial and haematopoietic fate decisions. Dll4 is highly enriched in arterial endothelium, promotes arterial specification and restricts hematopoietic development, consistent with high-intensity Notch signalling that maintain endothelial programmes (Duarte et al. 2004; Swift and Weinstein 2009). By contrast, Jag1 has been implicated in supporting definitive haematopoiesis without disrupting arterial identity (Robert-Moreno et al. 2008; Espin-Palazon et al. 2014; Monteiro et al. 2016), potentially by inducing a lower or more context-dependent Notch signalling output that permits HSPC emergence (Gama-Norton et al. 2015). Additionally, Jag1 plays a role in endothelial cells as a regulator of mechanotransduction pathways (Rodriguez et al. 2026), likely via Notch4 (Souilhol et al. 2022).

We have previously shown that *runx1* (a key regulator of HSPC fate) expression is reduced in *dll4* mutants (Bonkhofer et al. 2019), suggesting that *dll4* is required for the establishment or maintenance of hematopoietic gene expression in the hemogenic endothelium. Consistent with this observation, formation of HE derived from human pluripotent stem cells also requires DLL4 (Uenishi et al. 2018), indicating that there is a requirement for *dll4* for HE formation before HSPC emergence. Nevertheless, the mechanisms by which DLL4 regulates HE specification remain to be elucidated. Here, we identify Dll4 as an early regulator of hemogenic endothelial identity. Using single-cell transcriptomics, live imaging, and functional knockdown in zebrafish embryos, we demonstrate that *dll4* acts during HE specification, prior to any Jag1a-mediated effects on haematopoiesis. Mechanistically, Dll4 activity is required to fine-tune MAPK signalling, enabling endothelial cells to progress from pre-HE to HE and thereby commit cells towards a hematopoietic trajectory. *Jag1a* is then required sequentially, after the initial HE specification to promote HSPC emergence by mediating blood flow-dependent mechanical cues. Taken together, our findings reveal a temporally coordinated, cooperative activity of Dll4 and Jag1a ligands that integrate Notch signalling and flow-induced mechanical cues to establish HE identity and promote HSPC emergence. This uncovers an additional layer of regulatory complexity underlying the emergence of embryonic HSPCs.

## Results

### Dll4 is essential for hemogenic endothelium specification

Our previous work demonstrated a requirement for *dll4* in the expression of *runx1*, a key hemogenic endothelium marker and regulator of HSPC fate (Bonkhofer et al. 2019),supporting a role for *dll4* in haematopoiesis. Furthermore, studies indicated a dose-dependent requirement for Dll4 to induce arterial identity and restrict angiogenesis (Duarte et al. 2004; Leslie et al. 2007). To investigate whether this dose-dependent activity of Dll4 is required for HE formation and HSPC emergence, we injected a *dll4* splice morpholino oligonucleotide (*dll4 MO*)(Siekmann and Lawson 2007) into Tg(*kdrl:Hsa.HRAS-mCherry;Runx1P2:Citrine*) (Bonkhofer, 2019) embryos (here called *kdrl-cherry;runx1-citrine*) at increasing doses (7.5ng and 15ng) (Fig. S1A). *dll4 MO* activity was confirmed by PCR (Fig.S1B). As expected, this resulted in an increased number of ectopic vessels at 3 days post-fertilization (dpf) (Fig. S1A,C) with increasing doses of *dll4 MO*. Concomitantly, the number of *runx1-citrine*^+^ HSPCs in the CHT was also reduced in a dose-dependent manner (Fig.S1A,D). To determine whether *dll4* knockdown affects the broader haemogenic programme beyond runx1, we examined additional HE and early haematopoietic markers at 28 hpf., we first examined HE markers following *dll4* knockdown at 28 hpf. Expression of the HE markers *runx1*, *cmyb*, *gata2b*, *ikzf1,* and *gfi1aa* was significantly reduced in *dll4* morphants compared with wildtype embryos (Fig.1A,B, S1E,F). Importantly, while HE markers were significantly downregulated, the pan-endothelial marker *kdrl* and *flt4* (venous marker) were largely unaffected (Fig.S1F), suggesting that *dll4* loss primarily impairs HE specification rather than formation of the primary vasculature. In agreement with its role in establishing arterial identity, expression of the arterial markers *dlc* and *cldn5b* was downregulated (Fig. S1F).

**Figure 1.**
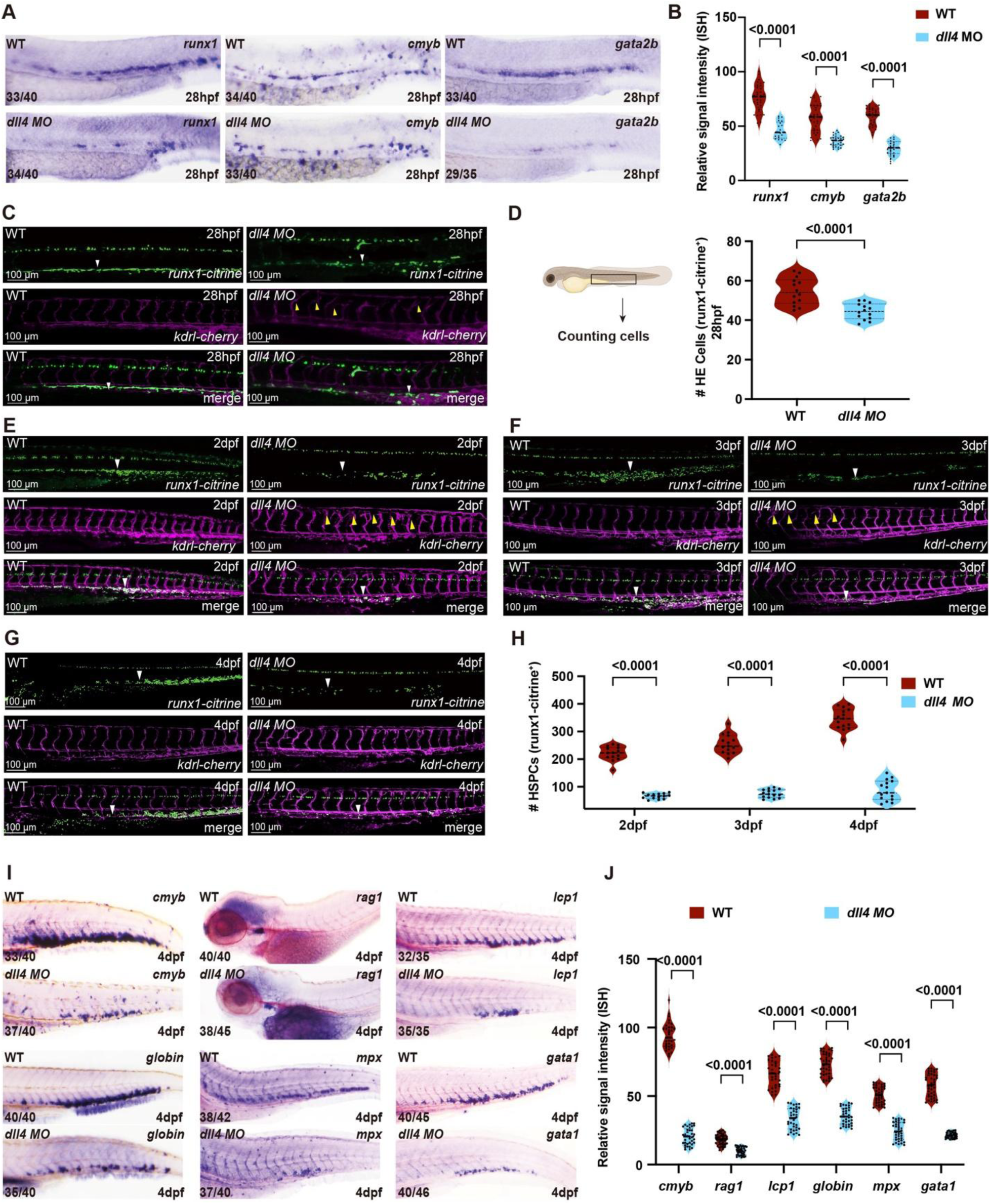
Dll4 is required for HE specification, HSPC emergence and differentiation. **(A**) Whole mount in situ hybridization (WISH) showing expression of *runx1*, *cmyb,* and *gata2b* at 28 hpf in WT and *dll4* morphants. Numbers in each panel indicate the number of embryos showing the representative phenotype over the total number analysed. **(B)** Quantification of WISH signal intensity shown in A. Each dot represents one embryo. Statistical significance was determined using unpaired T test. **(C)** Confocal imaging of *Tg*(*runx1:citrine);Tg(kdrl:mCherry*) embryos at 28 hpf. Green: runx1-citrine+cells; magnta – kdrl-Cherry+ endothelial cells. White arrowheads indicate HE cells, yellow arrowheads indicate ectopic sprouting of vessels. **(D)** Quantification of HE cells (*runx1:citrine⁺*) at 28 hpf shown in C. Each dot represents one embryo; samples were compared with an unpaired T-test. **(E–G)** Confocal imaging of *Tg*(*runx1:citrine);Tg(kdrl:mCherry*) embryos at 2 dpf (E), 3 dpf (F), and 4 dpf (G). Arrowheads indicate HSPCs. **(H)** Quantification of HSPCs (*runx1:citrine⁺*) at 2, 3, and 4 dpf shown in E–G. Each dot represents one embryo. Statistical significance was determined using unpaired T test. **(I)** WISH showing expression of hematopoietic lineage markers (*cmyb, rag1, lcp1, globin, mpx,* and *gata1*) at 4 dpf in WT and *dll4* morphants. Numbers indicate embryos showing the representative phenotype over total analysed. **(J)** Quantification of WISH signal intensity shown in I. Each dot represents one embryo. Statistical significance was determined using unpaired T test. Each data point represents one embryo. Data from three independent biological replicates were analysed. White arrowheads mark *runx1:citrine⁺* HE/HSPC cells, whereas yellow arrowheads indicate the hypersprouting phenotype in *dll4* morphants.

The reduced expression of HE markers could reflect either transcriptional downregulation or a loss of HE cells. To distinguish between these possibilities, we quantified the number of *runx1:citrine*⁺ HE cells in *dll4* morphants by confocal imaging in *kdrl-cherry;runx1-citrine* transgenics. The number of *runx1:citrine*^+^ HE cells was significantly decreased at 28hpf (Fig.1C,D), leading to a sustained reduction in *runx1-citrine*^+^ HSPCs from 2–4 dpf in *dll4* morphants (Fig.1 E-H). Concomitantly, aberrant vessel sprouting could be observed from 28hpf onwards (Fig.1 E-H). To confirm these phenotypes, we repeated this experiment in *dll4^sa9436^* homozygous mutants (here referred to as *dll4^-/-^*) in a *kdrl-cherry;runx1-citrine* background and confirmed the loss of HE at 28hpf, the severe reduction in *runx1-citrine^+^*HSPCs and the hypersprouting phenotype from 2-4dpf (Fig. S1I-K). Thus, we concluded that loss of *dll4* leads to a loss of HE and subsequently to a corresponding decrease in HSPCs produced in the embryo. Consistent with this decrease in HSPC numbers, expression of the T-cell marker (*rag1*) in the thymus and *cmyb* (HSPCs), *lcp1, mpx* (myeloid), and erythroid markers (*hbbe1, gata1*) in the CHT was downregulated in *dll4* morphants (Fig. 1 I,J) and *dll4^-/-^* mutants at 4dpf (Fig. S1L,M). Taken together, this indicated that *dll4* activity is required for HE specification to sustain the emergence of definitive HSPCs in zebrafish.

### Seeding of the CHT by endocardial- or dorsal aorta-derived HSPCs does not depend on Dll4 activity

Although reduced in number, HSPCs were still present in the CHT of *dll4* morphants at 3-4dpf (Fig. 1F-H). This was also observed in *dll4^-/-^* mutants (Fig. S1L,M), suggesting that there is an alternative source for the remaining HSPCs in the absence of *dll4*. We hypothesized that the recently identified endocardium-derived HSPCs (eHSPCs) (Bornhorst et al. 2024) might account for the remaining HSPCs seeding the CHT niche. To investigate this, we analysed the number of eHSPCs in the endocardium of *kdrl:BFP;CD41:GFP;runx1:mCherry* triple transgenic embryos at 48hpf, in the presence or absence of the *dll4 MO* (Fig. 2A,B). We detected a ∼2-fold decrease in the number of endocardial CD41:GFP^low+^ and *runx1*:mCherry^+^ HSPCs (Fig. 2A,B), indicating that these eHSPCs, similarly to those in the DA, require Dll4 activity. Next, we asked whether increased seeding of the CHT by eHSPCs could account for the remaining HSPCs observed in *dll4 MO* at 2dpf. Therefore, we used the photoconvertible protein Kaede together with a UAS-Gal4 system (Hatta, Tsujii, and Omura 2006) to perform lineage tracing of eHSPCs in the presence or absence of *dll4*. For that, we photoconverted the endocardium of *Tg(fli1a:Gal4*)*^ubs3^*;*Tg(UAS:Kaede*)*^rk8^*(Swift et al. 2014; Hatta, Tsujii, and Omura 2006) embryos (referred to as *fli1a:kaede*) transgenic embryos in control and *dll4* morphants at 48hpf (Fig. 2C,D) and counted photoconverted cells in the CHT at 4dpf (Fig. 2E). There was no difference in the number of eHSPCs seeding the CHT between control and *dll4* morphants, indicating that eHSPCs do not increase migration to the CHT. Photoconverting the DA at 48hpf yielded similar results (Fig. 2F-H). Taken together, these results indicated that neither eHSPCs nor late EHT events in the DA account for the remaining HSPCs observed in the CHT in the absence of *dll4*.

**Figure 2.**
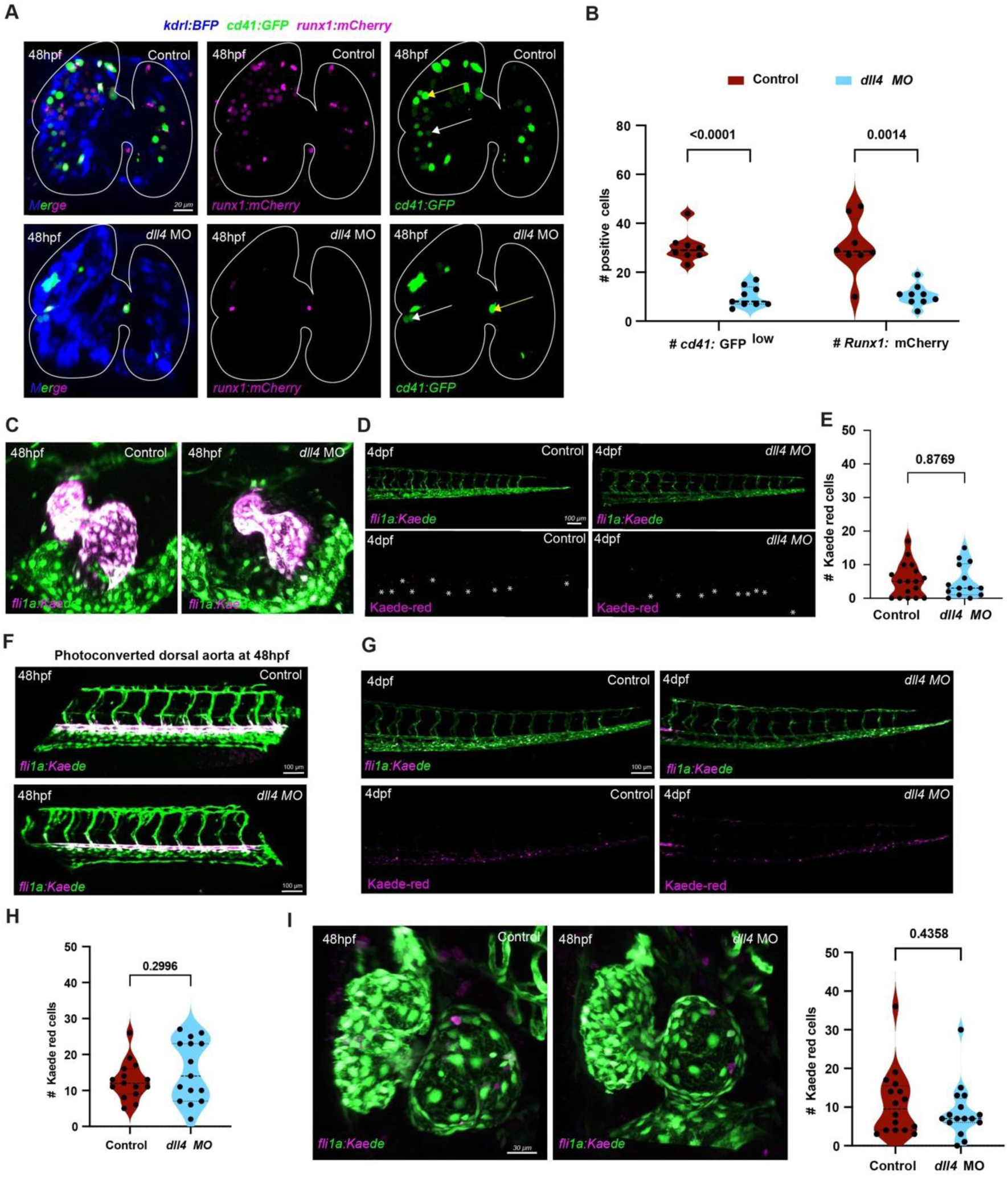
Endocardial HSPCs depend on Dll4 activity and do not compensate for the loss of dorsal aorta-derived HSPCs. **(A)** Confocal images of the heart in control and *dll4* MO at 48 hpf using Tg(*kdrl:BFP)* to label the endocardium*; Tg(cd41:GFP*) to label the HSPCs in GFP low expression; and *Tg(runx1:mCherry)* as an HSPC-only marker in embryos. Merged images and single-channel views of *runx1:mCherry* and *cd41:GFP* are shown. White dashed outlines mark the approximate cardiac region. White arrowheads indicate representative cd41:GFP^low^ (HSPC) and yellow arrowheads cd41:GFP^high^ (platelet) cells, respectively. **(B)** Quantification of *cd41:GFP*^low^ cells and *runx1:mCherry*-positive cells in the heart at 48 hpf. Each dot represents one embryo. **(C)** Confocal images of zebrafish hearts at 48 hpf showing cardiac endothelial cells labeled by fli1a:Kaede after photoconversion from green to red signal in control and *dll4* morphants. **(D)** Representative images of photoconverted heart-derived endothelial cells at 4 dpf in control and *dll4* morphant CHT regions. Upper panels show fli1a:Kaede (green/red); lower panels show Kaede-red (magenta). **(E)** Quantification of Kaede-red endocardial-derived cells following photoconversion being present in the CHT. Each dot represents one embryo. **(F)** Confocal images of zebrafish dorsal aorta at 48 hpf showing endothelial cells labeled by fli1a:Kaede after photoconversion from green to red signal in control and *dll4* morphants. Representative images show the region of photoconversion. **(G, H)** Representative images and quantification of photoconverted dorsal aorta-derived endothelial cells at 4 dpf in control and *dll4* morphants. Upper panels show fli1a:Kaede (green/red); lower panels show Kaede-red (magenta). **(I)** Confocal images and quantification of dorsal-aorta-derived Kaede-red endothelial cells in the heart at 4 dpf. Each dot represents one embryo. Scale bars: 100 µm unless otherwise indicated. Statistical comparisons and p-values are shown in the graphs. Statistical significance was determined using unpaired t-test.

### Dll4 and *Jag1a* cooperate to regulate hematopoietic stem and progenitor cell (HSPC) emergence and differentiation

To investigate whether other Notch ligands, expressed in endothelial cells or HE, could cooperate with Dll4 to regulate HSPC formation, we mined our previous data (Bonkhofer et al. 2019) for their expression in endothelial cells (Fig. S2A). The most prominent candidate expressed in endothelial cells was *jag1a*, which we and others have shown is required for HSPC formation(Espin-Palazon et al. 2014; Monteiro et al. 2016).Next, we first characterized whether these ligands cooperated to regulate vascular identity by comparing the expression of endothelial markers upon single or combined knockdown of *jag1a* and *dll4* using verified morpholinos (Siekmann and Lawson 2007; Yamamoto et al. 2010). Consistent with previous work (Robert-Moreno et al. 2008; Monteiro et al. 2016; Espin-Palazon et al. 2014),*jag1a* knockdown did not affect the expression of vascular identity markers at 28hpf. These included the pan-endothelial marker *kdrl*, venous markers (*flt4* and *ephb4a*), and arterial markers (*cldn5b* and *dlc*) (Fig. S2C,D). Loss of *dll4*, either alone or in combination with *jag1a MO*, led to increased *flt4* expression in tip cells (Fig. S2 C,D) and decreased arterial expression of *dlc* and *cldn5b* (Fig. S2 E,F), consistent with its role in establishing arterial identity (Ke et al. 2024). Confocal imaging of *kdrl:cherry* embryos demonstrated that vascular patterning was normal in control and *jag1a MO*, whereas *dll4 MO* and *jag1a+dll4* double *MO* embryos exhibited hypersprouting of intersegmental vessels at 4dpf (Fig.S2B). Altogether, these experiments confirmed that Dll4 (Siekmann and Lawson 2007), but not Jag1a, play a role in arterial development.

Next, we investigated whether Jag1a cooperated with Dll4 to specify HE. Hence, we assessed expression of HE markers at 28hpf following single or double MO knockdown of dll4 and jag1a (Fig. 3). Expression of *runx1*, *gata2b,* and *cmyb* (Fig.3 A, B) and *gfi1aa* and *ikaros* (Fig.S2G,H) were decreased in single *jag1a* and *dll4* morphants and further reduced in *jag1a+dll4* double morphants at 28hpf (Fig. 3A,B, Fig. S2G,H), indicating a defect in HE programming. Consistent with these transcriptional changes, confocal imaging of *runx1:citrine* embryos revealed a significant loss of *runx1:citrine⁺* HE cells at 28hpf in single *jag1a*- and *dll4* morphants, with a substantial reduction in combined *jag1a+dll4 MO* compared to either single knockdown (Fig. 3C,D). From 2 to 4 dpf, the number of *runx1:citrine⁺* HSPCs in the CHT remained substantially lower in both single *jag1a* and *dll4* morphants, with a more severe effect in the *jag1a*, *dll4* double morphant (Fig. 3E–H). Accordingly, expression of the HSPC marker *cmyb* in the CHT at 4 dpf was decreased in *dll4* or *jag1a* single morphants and almost completely abrogated in double *jag1a+dll4* morphants (Fig. 3I,J) compared to controls, indicating a severe loss of HSPCs. Consistent with this, expression of *gata1* (erythroid) and *lcp1* (myeloid) markers in the CHT and *rag1* (lymphoid) in the thymus were downregulated in single morphants and almost completely absent in double *jag1a+dll4* morphants at 4dpf (Fig. 3I,J). Taken together, these results demonstrate that Dll4 and Jag1a cooperate to promote HE specification and HSPC emergence.

**Figure 3.**
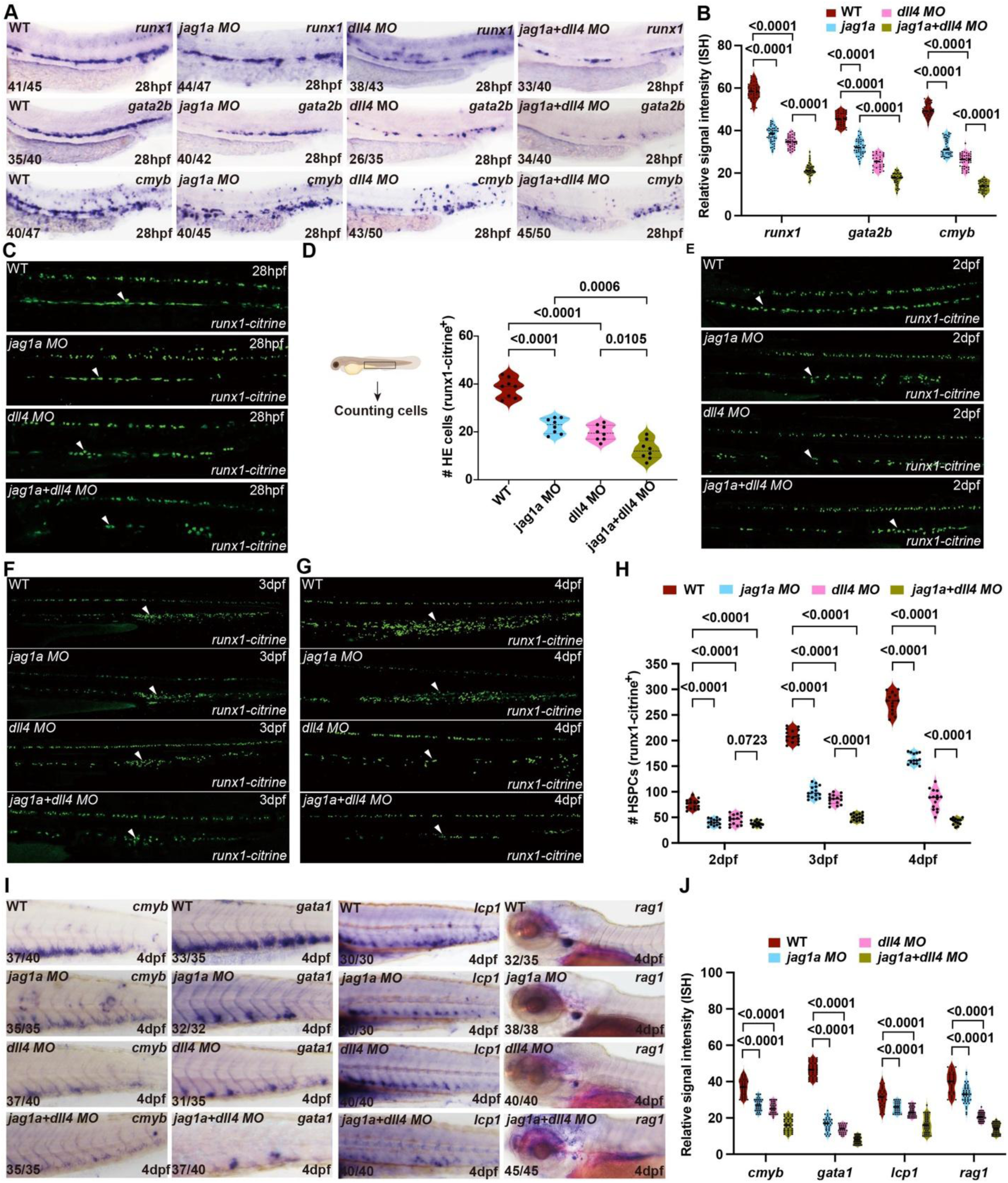
Cooperative *jag1a* and *dll4* activity regulates HSPC formation. **(A)** WISH showing expression of *runx1*, *gata2b,* and *cmyb* at 28 hpf in WT, *jag1a* morphants, *dll4* morphants and *jag1a + dll4* double morphants. Numbers indicate embryos showing the representative phenotype over the total number analysed. (**B)** Quantification of WISH signal intensity shown in A. Each dot represents one embryo. Statistical significance was determined using one-way ANOVA. **(C)** Confocal imaging of Tg(*runx1:citrine*) embryos at 28 hpf under the indicated conditions. Arrowheads indicate HE cells. **(D)** Quantification of HE cells (*runx1:citrine⁺*) at 28 hpf shown in C. Each dot represents one embryo (WT: n = 8; *jag1a MO* :n = 8; *dll4 MO*: n = 8; *jag1a+dll4* MO : n = 8); samples were compared with an unpaired T-test. **(E–G)** Confocal imaging of Tg(*runx1:citrine*) embryos at 2 dpf (E), 3 dpf (F), and 4 dpf (G) under the indicated conditions. Arrowheads indicate HSPCs. **(H)** Quantification of HSPCs (*runx1:citrine⁺*) at 2, 3, and 4 dpf shown in E–G. Each dot represents one embryo. Statistical significance was determined using one-way ANOVA. **(I)** WISH showing expression of hematopoietic lineage markers (*cmyb*, *gata1*, *lcp1,* and *rag1*) at 4 dpf under the indicated conditions. Numbers indicate embryos showing the representative phenotype over total analysed. **(J)** Quantification of WISH signal intensity shown in I. Each dot represents one embryo. Statistical significance was determined using one-way ANOVA.

### Dll4 activity regulates the transition from pre-HE to HE

Having established that both *dll4* and *jag1a* are required for HE specification at 28hpf, we further investigated whether these changes are detectable prior to that stage, at the onset of *runx1* expression in the HE (Wilkinson et al. 2009). Therefore, we analysed the expression of HE markers *runx1, gfi1aa,* and *gata2b* (Fig.4A,B) in single and double morpholino at 24hpf, Strikingly, *jag1a* morphants showed no changes in HE marker expression, while they were all severely reduced in *dll4*- or *jag1a+dll4* morphants (Fig. 4A,B). This indicates that, at the earliest stage of detectable HE marker expression, *dll4* is already required for HE gene expression, whereas *jag1a* is not. At 26 and 28hpf, *runx1* expression became progressively reduced in *jag1a* morphants, while the combined *jag1a*+*dll4* knockdown resulted in the strongest reduction of this HE marker (Fig. S2I,J). Thus, these data provide direct evidence that HE cells display distinct requirements for Notch ligand activity in endothelial cells between 24hpf and 28hpf. Although this developmental window is only 2–4h, it appears critical for defining whether haemogenic endothelium is specified. Taken together, these findings indicate that Dll4 activity is required earlier for HE specification, while Jag1a functions later, from ∼26hpf, cooperating with Dll4 to maintain HE programming and enable HSPC emergence.

**Figure 4.**
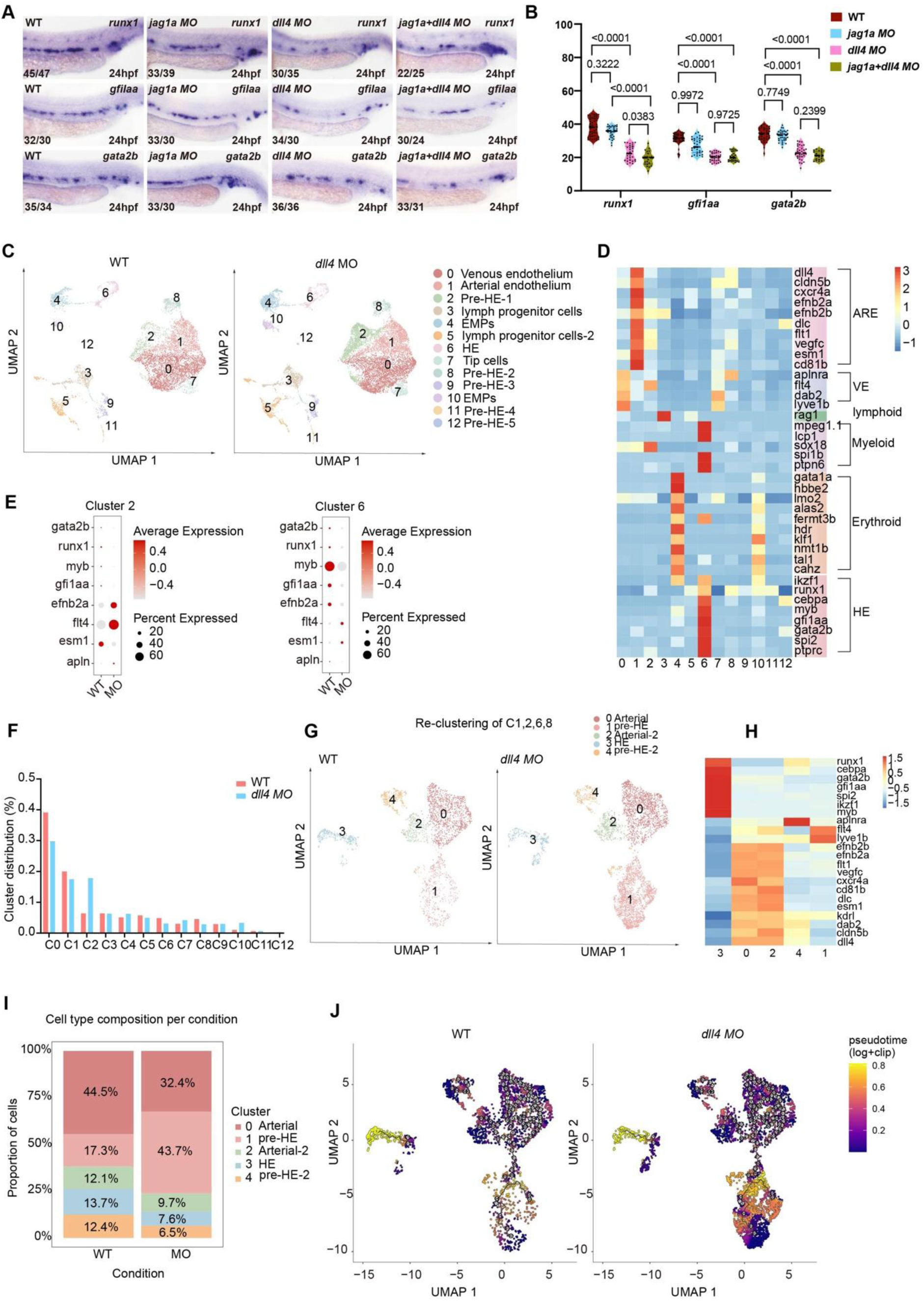
Dll4 is required early to enable programming of the hemogenic endothelium from a pre-hemogenic endothelium transcriptional state. (A) WISH for *runx1*, *gfi1aa*, and *gata2b* in WT, *jag1a* MO, *dll4* MO, and *jag1a*, *dll4* double MO at 24 hpf. Numbers indicate embryos showing the representative phenotype over the total number analysed. (B) Quantification of WISH signal intensity shown in A. Each dot represents one embryo. Statistical significance was determined using one-way ANOVA. (C) UMAP projections of single-cell RNA-seq data from WT and *dll4* MO showing annotated endothelial and hematopoietic populations, including arterial endothelium, venous endothelium, pre-hemogenic endothelium (pre-HE), hemogenic endothelium (HE), lymphoid progenitors, erythroid progenitors, myeloid progenitors, and related subsets. Cluster identities are indicated. (D) Heatmap showing scaled expression of selected marker genes across annotated clusters. The colour scale represents relative gene expression, with blue indicating lower expression, yellow indicating intermediate expression and red indicating higher expression.(E) Dot plots displaying average expression and percentage of expressing cells for selected markers in representative clusters (Cluster 2 and Cluster 6) in WT and *dll4* MO.Dot colour represents the scaled average expression of each gene, while dot size represents the percentage of cells within each cluster and condition with detectable expression of that gene.(F) Bar plot showing the proportion of cells in each cluster (C0–C12) in WT and *dll4* MO. (G) Heatmap of selected genes across re-clustered arterial endothelial and pre-HE and HE populations. (H) UMAP projections showing re-clustering of arterial endothelial and pre-HE/HE populations in WT and *dll4* MO.(I) Proportion of WT and *dll4* MO. (J) Pseudotime trajectory analysis of WT and *dll4* MO displayed on UMAP. Cluster 0 was used as the root for pseudotime trajectory.

To investigate how Dll4 contributed to HE programming, we isolated 24hpf control and *dll4 MO*-injected kdrl:GFP^+^ (Jin et al. 2005) endothelial cells for single-cell RNA sequencing (Fig. 4C). 18,913 cells (7,365 wt and 11,548 *dll4 MO*) passed quality control and were separated in 13 clusters (Fig. 4C) according to expression of key markers (Fig. 4D, Fig. S3A) and based on a previously used nomenclature (Ghersi et al. 2023). We defined venous endothelium (C0) and tip cells (C7), enriched in *flt4*, *dab2*, *lyve1b*; arterial endothelium (C1, enriched in *dll4*, *efnb2a*, *cldn5b*, cd81b), erythro-myeloid progenitors (C4, C10, enriched in *lmo2*, *gata1*, *klf1*), lymphoid progenitor clusters (C3, C5) and five pre-HE clusters (C2, C8, C9, C11, C12), enriched in *runx1* and low levels of various arterial or venous markers (Fig. 4D, S3B), suggesting heterogeneity in pre-HE populations. Among these, C2 showed stronger arterial/endothelial marker expression, suggesting an arterial-primed pre-HE identity. In contrast, the other pre-HE clusters showed variable enrichment of HE-associated and early haematopoietic markers, indicating heterogeneity within the pre-HE compartments rather than a single uniform cell state. We also identified one HE cluster (C6), highly enriched in markers including *runx1*, *cebpa*, *gata2b*, and *gfi1aa* (Fig. 4D). The main pre-HE cluster C2 and the HE cluster C6 express HE markers (*gata2b*, *runx1*, *myb*, *gfi1aa*), which were downregulated in *dll4* morphants (Fig. 4E). The reduction of HE markers in C2 may therefore reflect a selective loss or failure to maintain this early pre-HE population in *dll4* morphants. By contrast, *efnb2a* (arterial marker) was upregulated in C2 and some venous/tip cell endothelial markers (C2 - *flt4*, *apln*; C6 – *flt4*, *esm1*) were upregulated in the *dll4* morphants (Fig. 4E). This indicates that Dll4 is required not only to promote pro-haematopoietic gene expression but also for dampening the expression of arterial/pro-angiogenic genes (Helker et al. 2020; Hogan et al. 2009; Qiu et al. 2025). Consistent with this, the proportion of HE cells in C6 was decreased in *dll4* morphants (Fig. 4F). Strikingly, the proportion of pre-HE cells in cluster C2 was increased in *dll4* morphants by over ∼2-fold compared to the control (Fig. 4F, S3C). Together, these data suggest that loss of *dll4* impairs the progression from pre-HE to HE states, leading to an accumulation of pre-HE-like cells.

To investigate the dynamics of these changes in more detail, we re-clustered the arterial endothelial cells, pre-HE, and HE cells. We identified five subpopulations: two arterial (ARE-1 and -2, C0,C2), two pre-HE (C1,C4), and one HE (C3) cluster (Fig. 4G, H). Compared to the control cells, *dll4* morphants show an accumulation of cells at the less differentiated pre-HE1 state, resulting in fewer cells reaching the later HE cell stage (Fig. 4I). Next, we used Monocle 3 (Vocking and Famulski 2023) to perform pseudotime analysis on the re-clustered populations, with the ARE-1 population set as the origin (Fig. 4J). To further investigate transcriptional dynamics during HE progression, gene expression patterns were examined along pseudotime trajectories in wildtype and *dll4* morphants. In wildtype embryos, expression of *cmyb* gradually increased along pseudotime, consistent with a differentiation trajectory from ARE to HE. By contrast, *cmyb* expression remained reduced in *dll4 MO*, indicating impaired hematopoietic specification (Fig. S3D). The Notch ligand *jag1a* displayed low and transient expression along the trajectory in both conditions, consistent with its limited involvement during early HE specification (Fig. S3D). Notably, the MAPK pathway component *mapkapk2a* showed sustained low expression along pseudotime in wildtype embryos, whereas its expression was elevated in later stages in *dll4* morphants (Fig. S3D). We concluded that loss of *dll4* alters the normal progression from pre-HE to HE, causing cells to accumulate in a pre-HE state.

### Upregulated MAPK signalling drives the accumulation of pre-hemogenic endothelium in the absence of *dll4*

To identify signalling pathways associated with transcriptional changes in the major pre-HE (C2) and HE (C6) subpopulations (Fig. 4C), we first performed differential expression analysis in C2 (Fig. 5A). In this cluster, multiple MAPK-associated genes, including *mapkapk2a*, *rps6kal*, *prkd2,* and *insrb*, were significantly upregulated (Fig. 5B). A substantial number of MAPK pathway components, including *mapk1*, *mapk8*, *stat3,* and *mapkapk2a*, also showed increased expression in C6 (Fig. 5C). Accordingly, KEGG pathway enrichment analysis on the individual clusters (Fig. 5D,E, Fig. S4A-B) revealed significant enrichment of pathways related to calcium signalling, apelin signalling, and MAPK signalling in pre-HE (C2) (Fig. 5D). Similarly, KEGG pathway analysis of upregulated genes in C6 identified MAPK signalling as the most significantly enriched pathway (Fig. 5E). Together, these analyses identify MAPK signalling as a recurrent and enriched pathway in hemogenic endothelial subclusters of *dll4* morphants.

**Figure 5.**
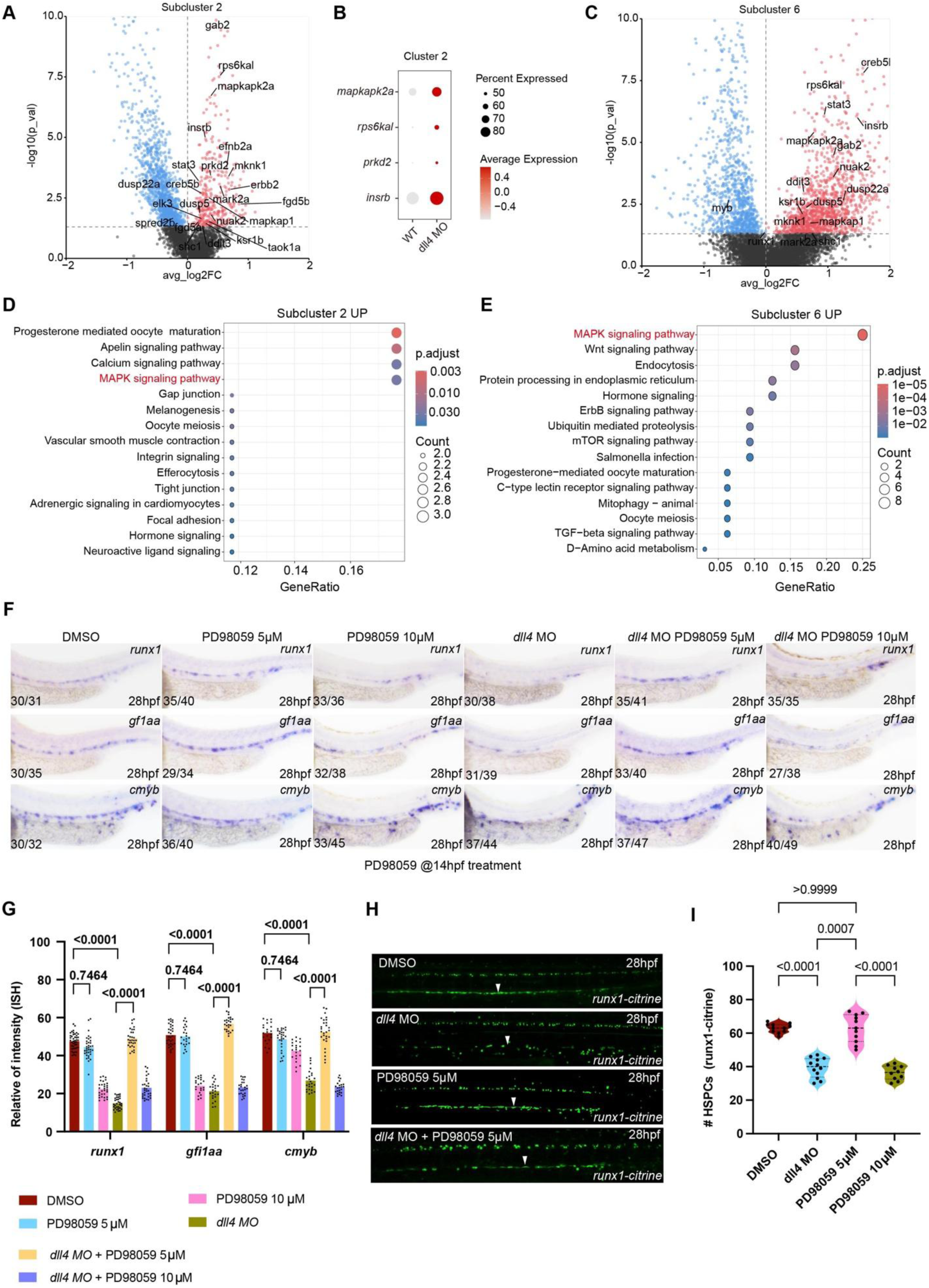
MAPK signalling acts downstream of dll4 to enable the transition from pre-HE to HE. **(A, C)** Volcano plots showing differential gene expression in cluster 2 and cluster 6 following *dll4* knockdown. **(B)** Dot plot showing expression of representative MAPK pathway genes across condition. Dot size represents the percentage of expressing cells and color indicates average expression. **(D, E)** KEGG pathway enrichment analysis of upregulated genes in cluster 2 (D) and cluster 6 (E). MAPK signalling pathway is significantly enriched among upregulated genes. Dot size indicates gene count and color represents adjusted p-values. **(F)** Whole-mount in situ hybridization (WISH) showing expression of *runx1, gf1aa,* and *cmyb* at 28 hpf under the indicated conditions. MEK inhibition (PD98059) partially rescues the loss of HE markers caused by *dll4* knockdown. Representative images are shown. **(G)** Quantification of WISH signal intensity for *runx1, gf1aa,* and *cmyb*. Each dot represents one embryo. Statistical significance was determined using one-way ANOVA. **(H)** Confocal images of Tg(*runx1: citrine*) embryos at 28 hpf showing HSPCs under the indicated conditions. Arrowheads indicate representative HSPCs. **(I)** Quantification of *runx1: citrine⁺* HSPCs per embryo. Each point represents one embryo. Statistical significance was determined using one way ANOVA.

Overactivation of the MAPK pathway represses the HE fate (Zhang et al. 2014). Therefore, we hypothesized that reducing the activity of the MAPK pathway in *dll4* morphants would rescue the block in pre-HE to HE transition. To test this, embryos were treated with 10 μM of the MEK inhibitor PD98059 from 14-28hpf to inhibit MAPK activity (Zhang et al. 2014). At this concentration, expression of the HE marker genes *runx1, gfi1aa,* and *cmyb* were downregulated compared to the DMSO control (Fig. 5F,G) as previously described (Zhang et al. 2014); at a lower concentration of PD98059 (5μM), no significant changes in the expression of these genes were observed after treatment (Fig. 5F,G). Notably, treatment with 5μM PD98059 efficiently rescued the expression of all three HE genes (*runx1*, *gfi1aa*, *cmyb*) at 28hpf (Fig. 5F,G), while the higher 10μM PD98059 failed to rescue HE marker expression. Consistent with these findings, analysis of *runx1:citrine* embryos at 28hpf revealed a significant reduction in *runx1-citrine+* HE cells in *dll4 MO*, rescued by combined treatment with 5μM, but not 10μM PD98059 (Fig. 5H,I). Collectively, these results demonstrate that *dll4* is a key modulator of the MAPK signalling pathway, fine-tuning its levels by regulating expression of its components during the transition from (arterial) endothelial cells to the hemogenic fate, thus enabling the emergence of HSPCs.

### Jag1a is a mechanosensitive gene mediated via blood flow to maintain the hemogenic cell fate

While we provide the first evidence that Dll4 is required during the early stages of HE specification, Jag1a only appeared to contribute to expression of HE genes from 26hpf, after the onset of blood circulation. Blood flow is a conserved regulator of HSPC development (North et al. 2009) and is required to stabilize hematopoietic programming (Wang et al. 2011), and emerging HSPCs tend to localize to areas of low or oscillatory shear stress (OSS) in the embryonic DA (Cui et al. 2022). Interestingly, murine *Jag1* (and to a lesser extent, *Dll4*) is induced by OSS in coronary arteries, driving pro-atherogenic signalling in the endothelium (Souilhol et al. 2022). Therefore, we hypothesized that during embryonic development, *jag1a* is induced by blood flow in the dorsal aorta to drive HSPC maintenance.

To investigate this, we injected the *tnnt2a MO* (Wang et al. 2011) to reduce (0.14ng) or block (1.4ng) the heartbeat and blood flow (Campinho et al. 2020) (Fig. 6A). As expected, reducing blood flow did not affect HE marker expression, while blocking led to decreased expression of *cmyb*, *gfi1aa*, and *gata2b* at 30hpf (Fig. S5A,B). At 4dpf, output from the HE-derived HSPCs was reduced in a dose-dependent manner: expression of HSPC marker *cmyb* and erythroid (*hbbe1^+^*, *gata1^+^*) in the CHT and lymphoid (*rag1^+^*) progeny in the thymus were all reduced, more severely in the absence of blood flow than when blood flow was reduced (Fig. S5C,D). Next, we assessed *jag1a* and *dll4* expression by qPCR in *tnnt2a* morphants at lower and higher *MO* doses (Fig. 6B,C). Reducing or blocking blood flow resulted in an approximately 50% reduction in *jag1a* expression (Fig. 6B), while expression of *dll4* was only affected when blood flow was completely blocked (Fig. 6C). We confirmed this effect on *dll4* expression by using nifedipine at increasing concentrations to induce partial or complete loss of circulation (Gierten et al. 2020) (Fig. S5E-G). Increasing blood flow with BF170 hydrochloride (Liu et al. 2024) significantly increased *jag1a* but not *dll4* expression (Fig. 6D-F). Taken together, these experiments suggest that *jag1a* is mechanosensitive and exhibits greater responsiveness to altered blood flow than *dll4*.

**Figure 6.**
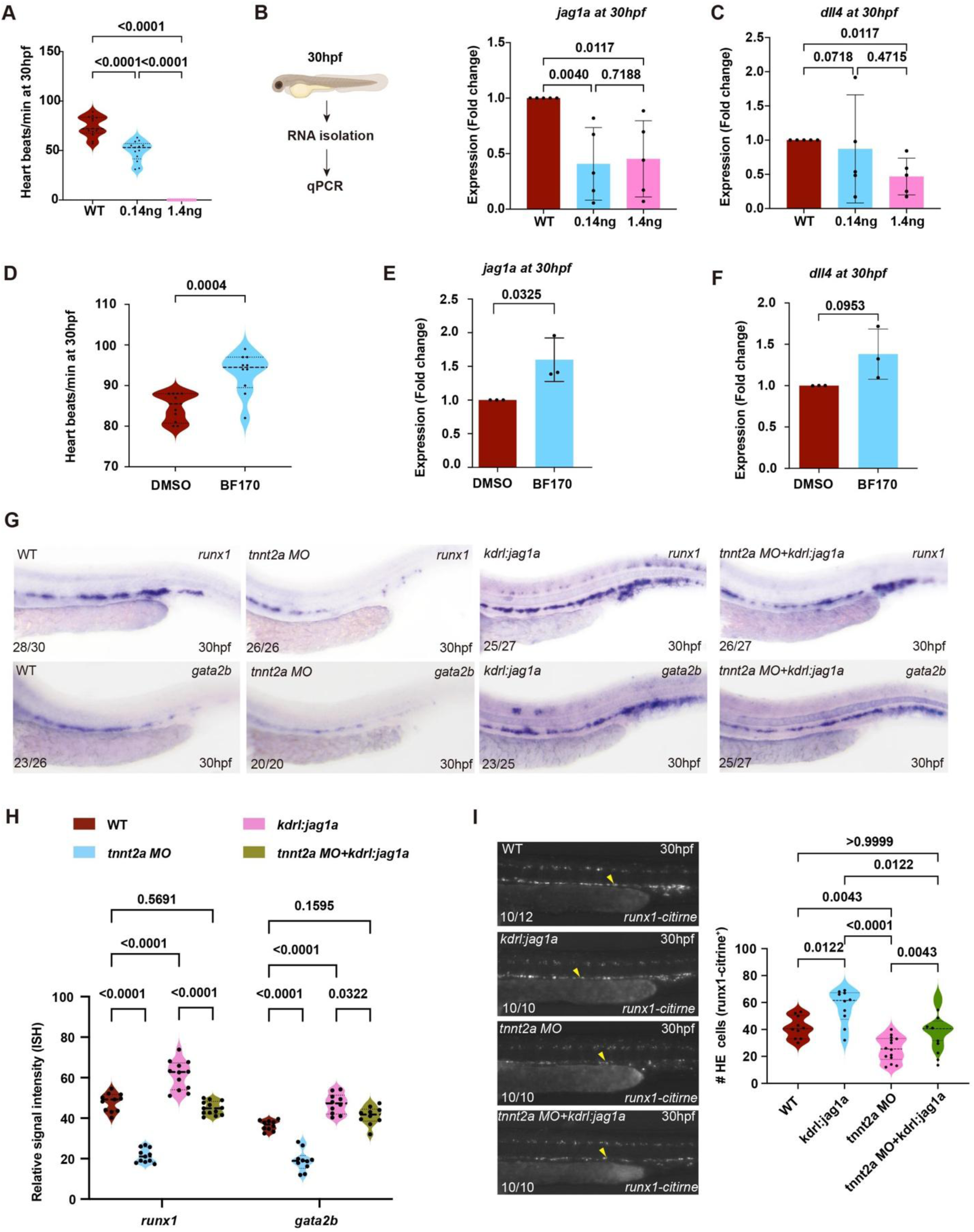
*jag1a* expression is regulated by blood flow and is required for HSPC formation from hemogenic endothelium. **(A)** dot represents one embryo (WT: n = 13; 0.14 ng: n = 13; 1.4 ng: n = 12). **(B, C)**Schematic of experimental design and quantification of heart rate at 30 hpf following *tnnt2a* knockdown. Each, qPCR analysis of *jag1a and dll4* expression at 30 hpf under the indicated conditions. Expression is shown as fold change relative to WT. Each dot represents one biological replicate (n = 5). Statistical significance was determined using unpaired two-tailed Student’s t-tests for pairwise comparisons. **(D)** Quantification of heart rate at 30 hpf following treatment with the Notch inhibitor BF170. Each dot represents one embryo (DMSO: n = 10; BF170: n = 11). Statistical significance was determined using an unpaired two-tailed Student’s t-test. **(E, F)** PCR analysis of *jag1a* (E) and *dll4* (F) expression at 30 hpf following BF170 treatment. Each dot represents one biological replicate (n = 3). Statistical significance was determined using an unpaired two-tailed Student’s t-test. **(G)** WISH showing expression of *runx1* and *gata2b* at 30 hpf in WT, *tnnt2a* morphants, *kdrl: jag1a* embryos and *tnnt2a* MO + *kdrl: jag1a*. Numbers indicate embryos showing the representative phenotype over total analysed. **(H)** Quantification of WISH signal intensity shown in G. Each dot represents one embryo. Statistical significance was determined using one-way ANOVA. **(I)** Living imaging of Tg (*runx1: citrine*) embryos at 30 hpf under the indicated conditions. Arrowheads indicate HE cells. Numbers indicate embryos analysed; Quantification of HE cells (*runx1: citrine⁺*) shown in I. Each dot represents one embryo (WT: n = 10; *kdrl: jag1a*: n = 10; *tnnt2a* MO: n = 14; double condition: n = 10). Statistical significance was determined using one way ANOVA.

To determine whether Jag1a is mediated via blood flow activity to maintain HE gene expression, we blocked circulation with the *tnnt2a MO* (1.4ng, Fig. 6E). Expression of the HE markers *runx1* and *gfi1aa* were downregulated at 30hpf (Fig. 6G,H). Endothelial-specific overexpression of *jag1a* under the *kdrl* promoter *(Monteiro et al. 2016)* rescued expression of both markers (Fig. 6G,H). Consistent with this, *jag1a* overexpression also rescued the loss of *runx1*:*Citrine*^+^ HE cells in the DA of *runx1-citrine* transgenic embryos at 30hpf (Fig. 6G). Taken together, these experiments indicate that Jag1a activity mediates mechanical cues from blood flow to ensure the expression of hematopoietic genes in HE and enable HSPC emergence.

## Discussion

Our study supports a model of Notch ligand function during zebrafish hemogenic specification, in which Dll4 acts early to enable entry into the hemogenic programme, whereas Jag1a acts later, after the onset of circulation, to maintain hemogenic identity and support downstream HSPC output. This framework extends the conventional view of Notch as a broadly permissive regulator of arterial and hemogenic fate by showing that distinct ligands contribute at different developmental phases and through mechanistically separable processes. Rather than acting as interchangeable activators of a common pathway, Dll4 and Jag1a appear to define successive regulatory modules that first establish, and then maintain, hemogenic competence. This logic is consistent with the idea that HSPC emergence is not controlled by a single specification event but instead requires progressive transition through intermediate endothelial states that remain sensitive to both intrinsic signalling and extrinsic biomechanical inputs (Horton et al. 2021; Ottersbach 2019).

Our findings are consistent with those of previous studies, indicating that hematopoietic cells in zebrafish originate from arterial endothelial cells and that the arterial environment is an indispensable prerequisite for subsequent hematopoietic differentiation (Bonkhofer et al. 2019). However, our data further suggest that the role of Dll4 in hematopoietic development is not limited to arterial patterning and instead it promotes the transition from pre-HE to HE by playing a broader role; it prevents expression of pro-angiogenic genes while favouring the expression of hemogenic genes in the nascent HE. This interpretation is more informative because it implies that Dll4 is required not only to induce specific genes, but also to tune the signalling environment in which endothelial cells become competent to progress along the hemogenic trajectory (Fig. 4).

In *dll4* deficient hemogenic endothelium, we observe an enrichment for MAPK signalling pathway genes (e.g. *mapk1, mapk8* and *mapkapk2a*). We show that partial MEK inhibition rescues hemogenic defects in *dll4* morphants, whereas stronger MEK inhibition suppresses hemogenic gene expression even in controls. These findings suggest a functional interaction between Dll4-dependent Notch signalling and the MAPK pathway during haemogenic specification to keep endothelial cells within a level of MAPK signalling that is permissive for hemogenic progression, a model supported by prior work indicating that modulation of the MAPK pathway enables establishment of the HE (Zhang et al. 2014). Taken together, these observations argue for a threshold-dependent model in which MAPK/ERK signalling must be neither too high nor too low but instead maintained within a narrow permissive corridor for pre-HE cells to progress to HE. The presence of *mapkapk2a* among the dysregulated genes raises the possibility that *dll4* loss may affect MAPK signalling beyond the canonical MEK/ERK pathway, particularly pathways involved in cytoskeletal remodelling and cell migration. The ERK pathway is known to regulate membrane proteins and cytoskeletal proteins (Roux and Blenis 2004). Given that EHT requires profound changes in cell shape, polarity and adhesion, it is plausible that MAPK dysregulation affects both hemogenic transcriptional identity and the morphogenetic execution of the transition (Kissa and Herbomel 2010; Boisset et al. 2010), although direct evidence for this remains to be established. Our current data therefore support a model of Dll4*-*dependent fine-tuning of MAPK signalling.

We also address the origin of residual HSPCs that persist after *dll4* knockdown; recent studies have shown that the zebrafish endocardium can contribute to embryonic hematopoiesis (Bornhorst et al. 2024), and a plausible explanation for the residual HSPC population in the CHT in the absence of *dll4* was compensatory endocardial seeding of the CHT (Bornhorst et al. 2024). However, we found that these eHSPCs also depend on Dll4 activity. Our photoconversion-based lineage tracing does not support increased endocardial contribution under *dll4* knockdown, nor do we observe evidence for enhanced late aortic seeding at 48hpf. These data substantially narrow the space of possible compensatory mechanisms. Residual HSPCs may instead reflect incomplete penetrance of *dll4* depletion, with partial redundancy with other Notch ligand-dependent pathways that permit limited hemogenic progression. This is an important point, because it suggests that while Dll4 is required for robust hemogenic specification, the system may retain a degree of buffering capacity that prevents complete collapse of HSPC production under partial perturbation.

Our analyses further indicate that Jag1a, another Notch ligand, acts at a later stage, after circulation begins, and cooperates with Dll4 to establish embryonic definitive hematopoiesis. This timing is notable because blood flow has long been recognised as a conserved requirement for embryonic HSPC development (Wang et al. 2011). Against this background, our findings show Jag1a as an interface between haemodynamic forces and sustained hemogenic programming. We find that *jag1a* expression is more sensitive than *dll4* to reduced or absent circulation, and that endothelial-specific *jag1a* overexpression is sufficient to restore hemogenic gene expression and HE cell number under flow-impaired conditions (Fig. 6). These results support the view that Jag1a does not play a role during the initial phase of hematopoietic induction but rather contributes to the maintenance of hematopoietic identity following the initial specification of the hemogenic fate. This is consistent with other studies showing that Jag1-mediated Notch signalling can respond to hemodynamic context and shape endothelial heterogeneity (Loerakker et al. 2018). We propose that a later *jag1a*-dependent maintenance module may act downstream of biomechanical stimuli to preserve hemogenic fate once initiated (Souilhol et al. 2022). How blood flow regulates *jag1a* expression in this context remains to be determined, but evidence from other model systems points to flow-activated KLF2 or KLF4 as potential regulators (Mack et al. 2009; Moonen et al. 2022).

Overall, our findings support a model in which hemogenic specification is governed by a sequential regulatory logic: Dll4 acts early to restrain MAPK signalling and thereby open a permissive window for pre-HE to HE progression, while Jag1a acts later to maintain hemogenic programming in a flow-dependent manner. This model has broader implications beyond zebrafish development. In mammals, definitive HSPCs also emerge from arterial haemogenic endothelium, and Notch signalling is required for both arterial specification and the generation of haematopoietic progenitors from haemogenic endothelium (Uenishi et al. 2018; Thambyrajah and Bigas 2022). Moreover, recent work suggests that Jag1 can regulate HSC identity within intra-aortic haematopoietic clusters in the mouse embryo, supporting the idea that distinct Notch ligands may exert stage- or context-specific functions during haematopoietic development (Thambyrajah and Bigas 2022). One of the central obstacles in HSPC engineering is that generating generic endothelium is not sufficient; the challenge is to recreate a developmental trajectory that first acquires hemogenic competence and then stabilises it under appropriate environmental conditions. We provide a conceptual basis for staged differentiation strategies in which MAPK/ERK activity is carefully tuned during hemogenic induction and flow-mimicking chemical or mechanical signals are introduced later to support stable HSPC output.

## Material and Methods

### Animal maintenance and genotyping

Adult wildtype, *dll4^sa9436^* mutants, TgBAC(*runx1P2:Citrine*;*kdrl:mCherry*) double transgenic zebrafish (here referred to as *runx1:citrine;kdrl:cherry*)(Bonkhofer et al. 2019), *dll4^sa9436/+^*;*runx1:citrine*;*kdrl:Cherry,* Tg(*kdrl:BFP-CAAX)^mu293^*(Matsuoka et al. 2016), *Tg(Runx1:mCherry)(Tamplin et al. 2015)*, *Tg(fli1a:Gal4)^ubs4^* (Hatta, Tsujii, and Omura 2006) and *Tg(UAS:Kaede)^rk8^* (Swift et al. 2014) were maintained under standard conditions at 28.5°C with a 12h light/12h dark cycle. All procedures involving zebrafish were conducted in compliance with the UK Animals Scientific Procedures Act (ASPA) 1986, under project licence PP5005485. Genotyping of the *dll4*^sa9436^ allele was performed using a KASP-TF master mix (Biosearch Technologies, Cat#KBS-1050-121) targeting the custom sequence CGGCCACTACACCTGCAACCCAGATGGCCRGTTATCCTGTCTCCCTGGCT[G/A]GAAGGGGGAATACT GCGAAGAACGTAAGCAATCAGAGWTCACAATTTATT, following the manufacturer’s instructions (Bonkhofer et al. 2019).

### Morpholinos, DNA injections, and chemical inhibitors

Morpholino oligonucleotides (Gene Tools) designed to block *dll4 (dll4 MO: 5’-CGAATCTTACCTACAGGTAGATCCG-3’(Bonkhofer et al. 2019), jag1a (jag1a MO*: 5’-AAGCCAAACCCGCACATACCCGCAT-3’(Yamamoto et al. 2010), and *tnnt2a* (*tnnt2a MO*:5′-CATGTTTGCTCTGATCTGACACGCA-3′(Sehnert et al. 2002), were injected at the 1-2 cell stage, typically 1 nL per embryo, at the concentrations indicated in the figures. Overexpression of jag1a in endothelial cells to rescue the loss of blood flow was performed using a kdrl:jag1a-V5 construct as previously described(Monteiro et al. 2016).

To explore whether reducing the activity of the MAPK pathway could rescue the phenotype of dll4 morphants, we treated wildtype and dll4 *MO*-injected embryos with the selective MEK inhibitor PD98059 (HY-12028,Cambridge Bioscience) from 14hpf-30hpf at high (10μM, (Zhang et al. 2014) and low (5μM) concentration prior to *in situ* hybridization or live imaging of the *runx1:citrine* transgenic.

### Whole mount in situ hybridization

Whole-mount in situ hybridization (WISH) was performed as previously described(Jowett and Yan 1996), using established probes for key markers (Supplementary Table1). Following hybridization, embryos were imaged under standardized illumination and exposure on a Nikon SMN800N stereomicroscope equipped with a Nikon DS-Fi3 camera and NIS-Elements F software. Images were stored as .TIFF files and quantification of gene expression was performed as previously described, with signal intensity and area measured using ImageJ software (Dobrzycki et al. 2018) Corrected intensity values are displayed as scatter dot plots and statistical analyses performed using GraphPad Prism 10; statistical tests are described in figure legends.

### mRNA extraction, cDNA synthesis, and quantitative RT–PCR

Total RNA was isolated from the zebrafish embryos using the Zymo RNA Extraction Kit (Zymo, Cat# R2130). cDNA was synthesized from total RNA using a Superscript IV RT-PCR enzyme (Invitrogen, Cat# 12574026) following the manufacturer’s instructions. qRT–PCR was performed on a QuantStudio Real-Time PCR (Thermo Fisher Scientific) with *Power* SYBR™ Green PCR Master Mix (Applied Biosystems, Cat#4367659). The primers used for qRT–PCR are listed in Supplementary Table 2. Fold changes in gene expression were calculated using the 2^−ΔΔCt^ method (Livak and Schmittgen 2001), Expression levels were normalized to the housekeeping gene *rplp0*.

### Confocal and epifluorescence Imaging

Confocal imaging was performed using a Leica Stellaris 8 DLS confocal microscope equipped with o60× objectives and Zeiss 710 and 40x for heart and 20x for tail images, all information can be found in(Bornhorst et al. 2024). Embryos were mounted in 1% low melting agarose in E3 medium, covered in E3 medium with tricaine and imaged at room temperature. Z-stack images were acquired at 0.5 µm layer. To obtain a larger field of view, adjacent image tiles were captured with 10% overlap and seamlessly reconstructed using the LAS X stitching module (Leica Microsystems). Images were processed in LAS X to generate maximum intensity projections. Alternatively, we imaged uninjected and morpholino-injected *kdrl:cherry;runx1:citrine* embryos on a Zeiss SteREO Discovery.V12 stereomicroscope with a Zeiss Axio Cam MRm and Axio Vision software. Cell counting was performed manually using ImageJ. GraphPad Prism 10 software was used to generate scatterplots of cell counts and for statistical analysis; statistical tests are described in the figure legends.

Embryos, up to 4dpf, were embedded in 1% low-melting agarose supplemented with 0.2% Tricaine. Images of stopped hearts and the CHT were acquired using a Zeiss LSM710 confocal laser microscope at 20x magnification for the CHT and 40x magnification for the heart with a dipping lens, respectively. Maximal-intensity projections were uniformly generated using identical settings for all samples. Image processing and subsequent quantitative analyses were conducted using Fiji software tools (Schindelin et al. 2012) and Imaris (Bitplane, UK). Brightness and contrast were adjusted with Imaris and Fiji.

### Zebrafish embryo, cell dissociation, sorting and scRNA library preparation

Zebrafish embryos were dechorionated manually using fine forceps and overdosed with anesthesia at 28hpf. The embryos were first incubated in deyolking buffer (116mM NaCL, 2.9 mM KCl, 5mM HEPES, 1mM EDTA) to remove the yolk, followed by incubation in dissociation mix (68% TrypLE Express (Gibco, 12605036), 5mg collagenase (Merck, C9891-500MG) in 1 x HBSS (Gibco, 14025092)) and incubation at 30°C for 10 minutes whilst regularly pipetting to fully dissociate the cells. Enzymatic activity was stopped by addition of FBS (Sigma-Aldrich, Cat#A3059) at a final concentration of 12.5%. Cells were pelleted at 300xg for 5 min, washed twice (2.5mg/ml BSA & 1x HEPES in 1 x HBSS) and filtered through a 30 µm strainer (Sysmex, Cat# 04-0042-2316). Cell suspensions were sorted for GFP (kdrl-GFP) positive cells using a BD FACSAria™ Fusion into DMEM with 10% FBS. Following sorting, the cells were incubated at 28°C for 15 minutes to allow recovery prior to centrifugation at 300xg for 5 minutes and washing with cell suspension buffer () and pelleting at 500xg for 5 minutes. Single-cell libraries were prepared using the Illumina Single Cell 3′ RNA Prep, T20 kit following manufacturer’s instructions and the resulting libraries sequenced by Novogene UK (Cambridge, UK) on an Illumina NovaSeq X Plus Series with a paired end 150bp run.

### Single-cell data analysis

Single-cell RNA-seq data were processed using PIPseeker (Fluent Biosciences) and analysed in R (v4.3.1) using Seurat (v5.0.0) (Hao et al. 2024) and Monocle3 (Vocking and Famulski 2023). Quality control filtering, normalization, clustering, and UMAP visualization were performed using Seurat. Differential gene expression analysis was performed using Seurat. Marker genes between conditions were identified using the Wilcoxon rank-sum test implemented in FindMarkers, with p-values adjusted using the Benjamini– Hochberg method. Genes with an adjusted p-value < 0.05 were considered significant. Volcano plots were generated using ggplot2 to visualize log2 fold changes and adjusted p-values. Heatmaps were generated using Seurat (DoHeatmap) based on scaled expression values of selected genes. Trajectory reconstruction and pseudotime analysis were conducted using Monocle3 (Vocking and Famulski 2023).Gene ontology (GO) and pathway enrichment analyses were performed using the Database for Annotation, Visualization and Integrated Discovery (DAVID, v6.8)(Dennis et al. 2003). Kyoto Encyclopedia of Genes and Genomes (KEGG) pathway enrichment (Kanehisa and Goto 2000) was conducted using DAVID or cluster Profiler in R, and pathways with adjusted P < 0.05 were considered significant.

### Photoconversion of endothelial cells

*Tg(fli1a:Gal4*)*^ubs3^*;*Tg(UAS:Kaede*)*^rk8^* (Swift et al. 2014; Hatta, Tsujii, and Omura 2006) embryos in which the photoconvertible protein Kaede is specifically expressed in endothelial cells were used for photoconversion experiments as previously described (Bornhorst et al. 2024). Briefly, defined vascular regions, including the endocardium and the dorsal aorta (DA), were selected for photoconversion using a Zeiss LSM710 confocal microscope. Photoconversion was performed by five scans with a 405 nm laser at 100% power. Z-stacks of the Kaede-green and Kaede-red channels were acquired before and after photoconversion to confirm spatially restricted conversion. For photoconversion within the DA, a region spanning 10 intersegmental vessels (ISVs) was defined as the standard region of interest. Following photoconversion, embryos were recovered and maintained at 28 °C until subsequent imaging.

## Acknowledgements

We are grateful to the staff of the Biomedical Services Unit for fish husbandry and Claire Bryer and Kate Orr of Genomics Birmingham (University of Birmingham) for genomic sequencing. We acknowledge the support of the Flow Cytometry Technology Hub Facilities (Ferdus Sheik and Megan Butler) at the College of Medicine and Health, University of Birmingham, for providing access to equipment and technical expertise. This research was supported by the Department of Cancer and Genomic Sciences, University of Birmingham, a British Heart Foundation Project Grant to R.M. and B.E-W. (PG/22/11161), and an Add-on Fellowship of the Joachim Herz Foundation to D.B. F.G. acknowledges funding from the DFG (Projects SFB1348B12 and SF1450N04), the Cells-in-Motion Interfaculty Center, the Faculty of Medicine at the University of Münster (the Innovative Medical Research grant (I-GU122208)), the Natural Sciences and Engineering Research Council of Canada (NSERC) Discovery Grant (Project RGPIN-2025-06873), the Canadian Foundation for Innovation and Ontario Research Fund John R. Evans Leadership Fund (Project 46346), and the Department of Cell and Systems Biology at the University of Toronto.

