## Supplementary Figures for "Dll4 and Jag1a signalling act sequentially and cooperatively to drive hematopoietic stem cell fate specification"

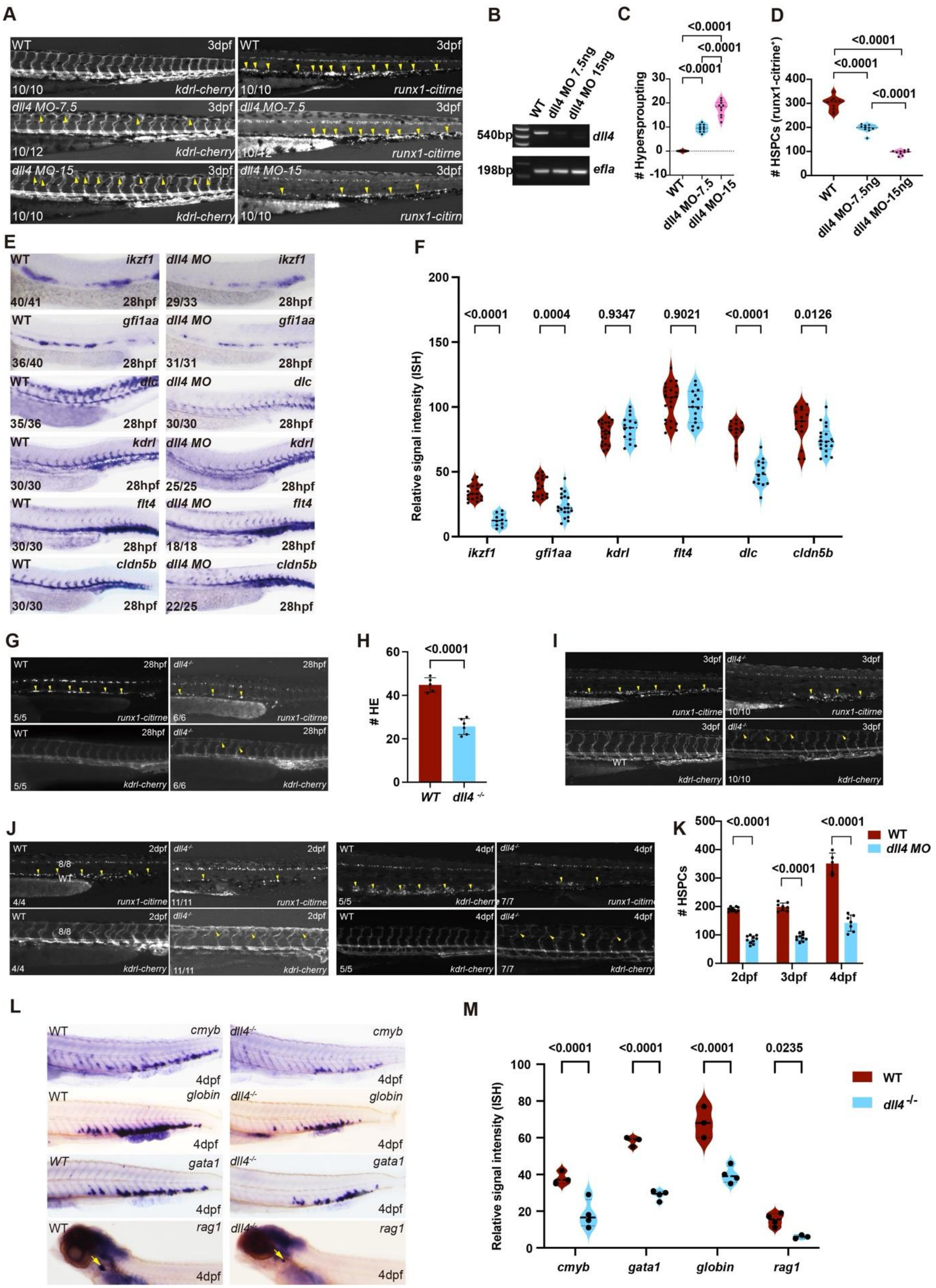

**Figure S1. Dose-dependent loss of *dll4* via MO knockdown or genetic ablation of *dll4* severely decreases HSPC formation in the dorsal aorta and lineage differentiation in the CHT. (A)** Confocal images of Tg(*runx1:citrine; kdrl:mCherry*) embryos at 3 dpf in WT embryos and embryos injected with 7.5 ng or 15 ng *dll4* MO. Lateral views of the trunk region are shown. Arrowheads indicate representative *runx1:citrine* positive cells. **(B)** PCR analysis of *dll4* transcripts in WT embryos and embryos injected with increasing doses of *dll4* MO. *ef1a* is shown as housekeeping gene. **(C,D)** Quantification of *runx1:citrine* positive HSPCs and hypersprouting at 3 dpf in WT and *dll4* MO embryos at the indicated doses. Each dot represents one embryo. Samples were compared with an unpaired T-test. **(E)** ISH at 28 for endothelial and hematopoietic markers, including *ikzf1, gfi1aa, dlc, kdrl, flt4,* and *cldn5b*, in WT and *dll4* MO embryos. Representative lateral trunk views are shown. **(F)** Quantification of relative ISH signal intensity for markers shown in (D). Each dot represents one embryo. Statistical significance was determined using unpaired T test. **(G-H)** Images of Tg(*runx1:citrine; kdrl:mCherry*) embryos at 28 hpf, 2dpf, 3 dpf, and 4 dpf in WT and *dll4* MO embryos. Arrowheads indicate representative *runx1:citrine*-positive cells and hypersprouting. Quantification of *runx1:citrine* positive HE cells at 28 hpf and HSPCs at 2, 3, and 4dpf in WT and *dll4* MO embryos. **(I,J)** Live imaging of 2 dpf, 3 dpf, and 4 dpf. Numbers in each panel indicate the number of embryos showing the representative phenotype over the total number analysed. **(K)** Quantification of WISH signal intensity. Each dot represents one embryo. Statistical significance was determined using unpaired T test. **(L)** ISH at 4dpf for lineage markers including *cmyb, gata1, globin*, and *rag1* in WT and *dll4* mutant embryos. Representative lateral views are shown. **(M)** Quantification of relative ISH signal intensity for markers. Statistical significance was determined using one way ANOVA or unpaired ttests as appropriate.

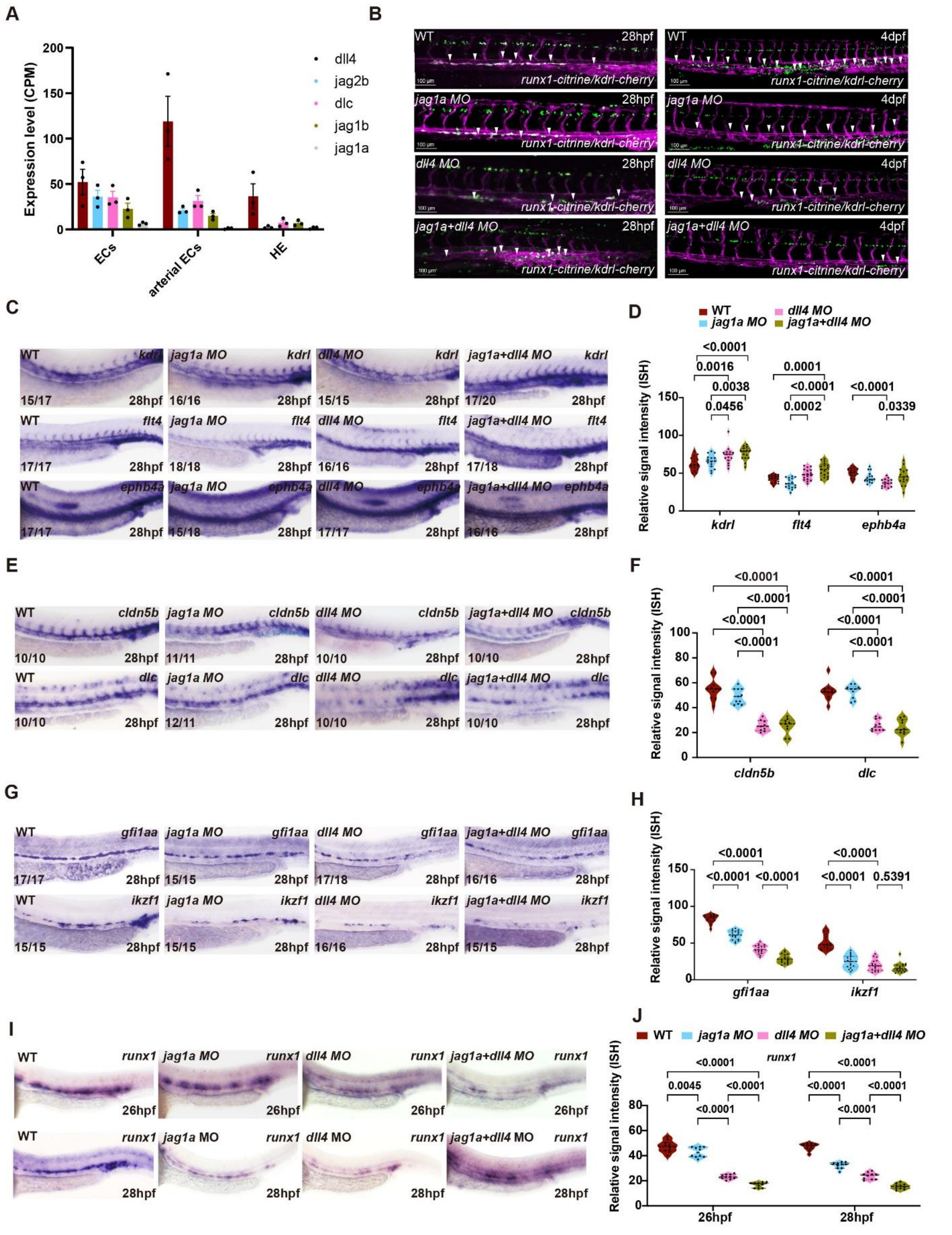


**Figure S2. Jag1a cooperates with Dll4 during hemogenic endothelium specification and HSPC emergence. (A)** RNA-seq expression of the Notch ligands *dll4*, *jag2b*, *dlc*, *jag1b* and *jag1a* in endothelial cells, arterial endothelial cells and haemogenic endothelium at 28hpf (data from Bonkhofer et al, 2019). Expression values are shown as counts per million reads (CPM).**(B)** Confocal imaging at 28hpf and 4dpf.The white arrow points to the HSPCs.**(C)** WISH showing arterial marker expression (*kdrl, flt4, ephb4a*) at 28 hpf in WT and *dll4-, jag1a-* and *dll4+jag1a* morphants. Numbers indicate embryos showing the representative phenotype over the total number analysed. **(D)** Quantification of WISH signal intensity shown in B. Each dot represents one embryo. Statistical significance was determined using one-way ANOVA **(E)** WISH showing expression of *cldn5b, dlc* at 28 hpf in WT and *dll4 MO, jag1a MO,* and *dll4+jag1a* morphants. Numbers indicate embryos analysed. **(F)** Quantification of WISH signal intensity shown in D. Each dot represents one embryo. Statistical significance was determined using one-way ANOVA. **(G)** WISH showing hemogenic endothelium (*gfi1aa, ikzf1*) expression at 28 hpf in WT and *dll4-, jag1a-* and *dll4+jag1a* morphants. **(H)** Quantification of WISH signal intensity shown in F. Statistical significance was determined using one-way ANOVA **(I)** WISH showing *runx1* expression at 26 hpf and 28 hpf in WT and *dll4-, jag1a-* and *dll4+jag1a* morphants. **(J)** Quantification of *runx1* signal intensity shown in H. Data are presented as mean ± s.d. Statistical significance was determined using one-way ANOVA.

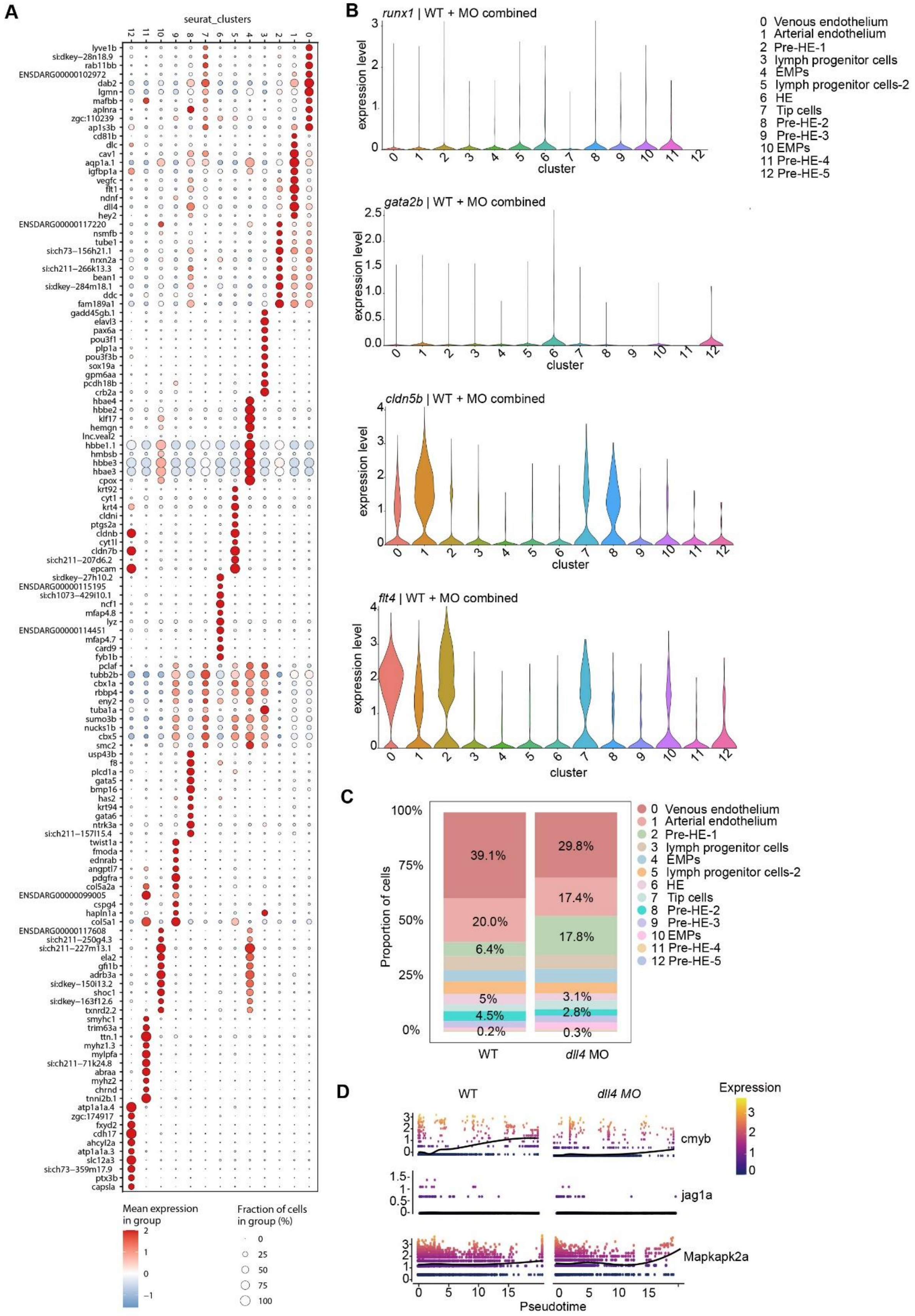


**Figure S3. Single-cell transcriptomic analysis of endothelial and hematopoietic populations in WT and *dll4* MO. (A)** Bubble plot showing the top 10 genes expression across identified cell clusters (0–12). Cells are grouped by cluster identity and colored according to normalized gene expression levels. **(B)** Expression of HE markers (*runx1*, *gata2b*), and vascular markers (*flt4*, *cldn5b*) across clusters. Each dot represents a single cell. **(C)** Stacked bar plot showing the proportion of annotated cell populations in WT and *dll4* MO, including venous endothelium, arterial endothelium, pre-HE, HE, pre-HE, lymphoid progenitors, EMPs, and HSPCs. **(D)** Expression dynamics of *cmyb*, *jag1a* and *mapkapk2a* along pseudotime in WT and *dll4* morphant cells. Each point represents an individual cell, coloured according to its gene expression level. The x-axis represents pseudotime and the y-axis represents gene expression. Black lines indicate fitted expression trends along pseudotime.

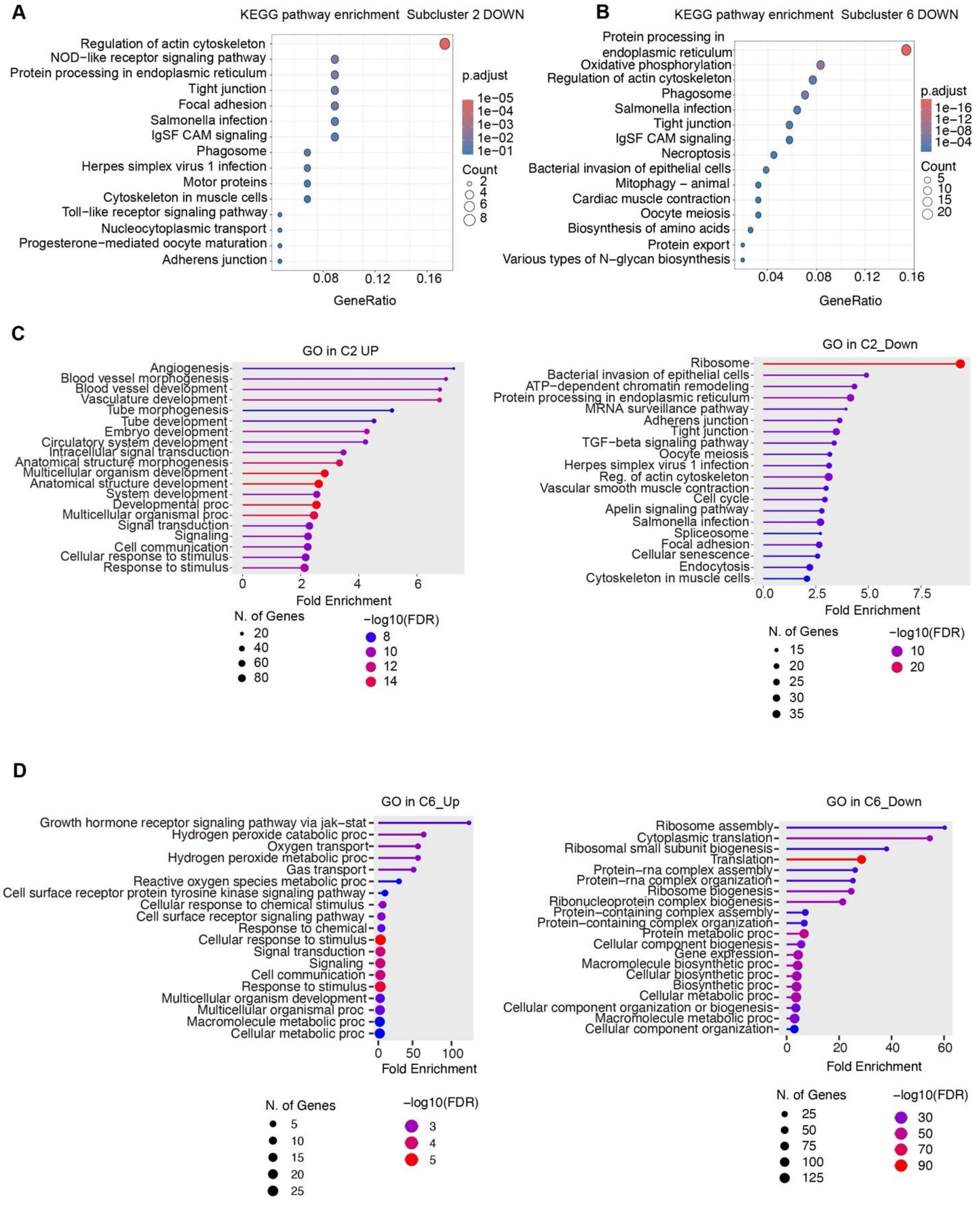


**Figure S4. Pathway enrichment analysis reveals distinct transcriptional changes in pre-HE and HE subclusters. (A, B)** KEGG pathway enrichment analysis of downregulated genes in subcluster 2 (A) and subcluster 6 (B). **(C)** Gene ontology (GO) enrichment analysis of upregulated (left) and downregulated (right) genes in subcluster 2. **(D)** Gene ontology (GO) enrichment analysis of upregulated (left) and downregulated (right) genes in subcluster 6.

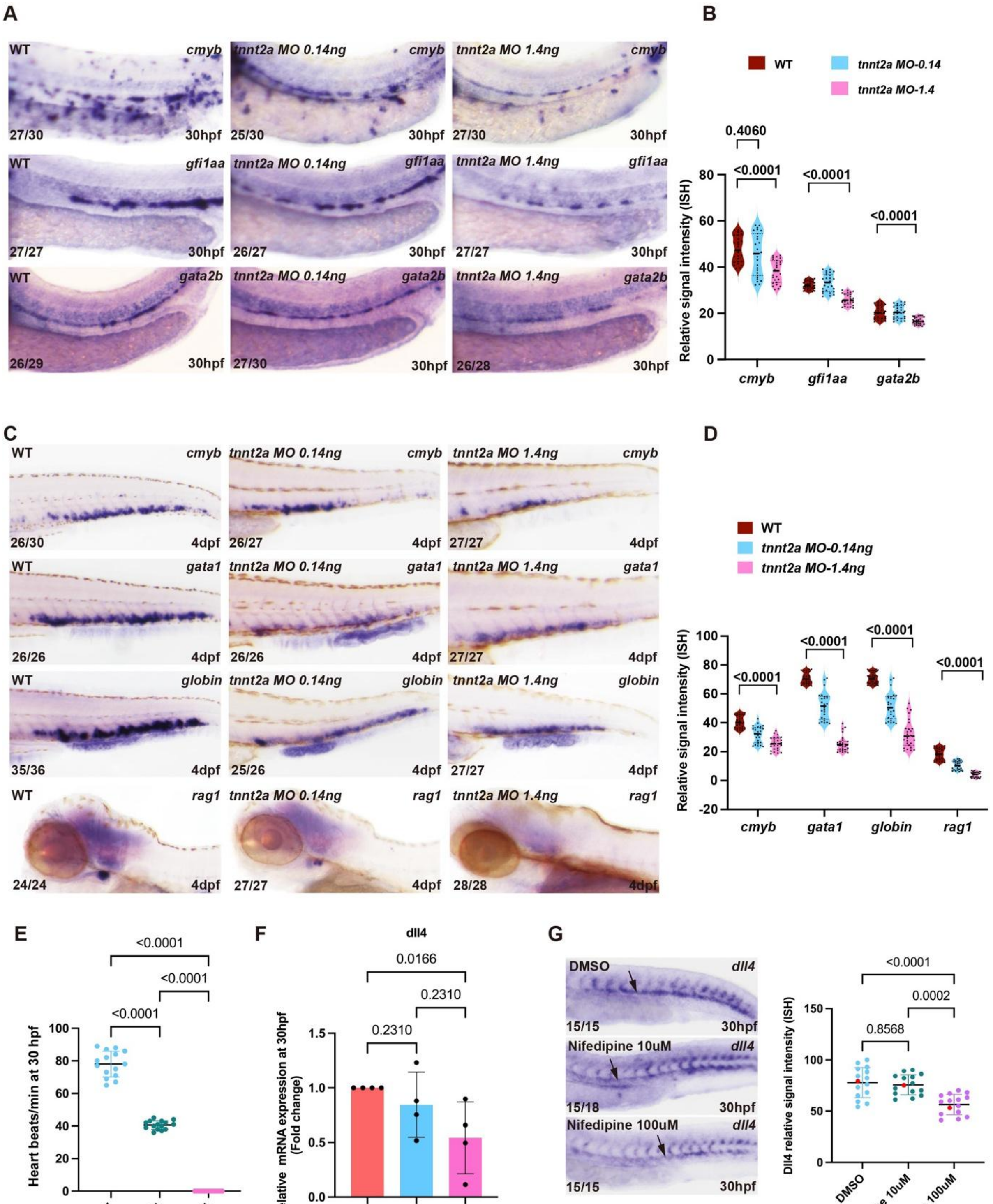


**Figure S5**. **Blood flow is required for maintenance of hemogenic endothelium gene expression and emergence of normal numbers of HSPCs. (A)** WISH showing expression of *cmyb, gfi1aa, gata2b,* and *runx1* at 30 hpf in WT and *tnnt2a* morphants. Numbers indicate embryos showing the representative phenotype over the total number analysed. **(B)** Quantification of WISH signal intensity shown in A. Each dot represents one embryo. Statistical significance was determined using one-way ANOVA. **(C)** WISH showing expression of hematopoietic lineage markers (*cmyb, gata1, globin,* and *rag1*) at 4 dpf in WT and *tnnt2a* morphants. Numbers indicate embryos showing the representative phenotype over the total number analysed. **(D)** Quantification of WISH signal intensity shown in C. Each dot represents one embryo. **(E)** Quantification of heart rate (beats per minute) at 30 hpf in WT embryos and embryos treated with nifedipine (10 µM and 100 µM). Each dot represents one embryo. **(F)** qPCR analysis of *dll4* mRNA expression at 30 hpf following nifedipine treatment. Expression is shown as fold change relative to DMSO control. Each dot represents one biological replicate. **(G)** Whole-mount in situ hybridization (WISH) showing dll4 expression at 30 hpf in DMSO-treated embryos and embryos treated with nifedipine (10 µM and 100 µM). Representative images are shown (left). Right, quantification of WISH signal intensity. Each dot represents one embryo. Arrows indicate regions of *dll4* expression. Statistical significance was determined using one-way ANOVA.
